# Principles of dengue virus evolvability derived from genotype-fitness maps in human and mosquito cells

**DOI:** 10.1101/2020.02.05.936195

**Authors:** Patrick T. Dolan, Shuhei Taguwa, Mauricio Aguilar Rangel, Ashley Acevedo, Tzachi Hagai, Raul Andino, Judith Frydman

## Abstract

Dengue virus (DENV), an arbovirus infecting over 100 million annually, cycles between human and mosquito hosts^1^. Examining how DENV adapts to such different host environments could uncover principles of arbovirus transmission and emergence. Here we combine sequential passaging and ultra-deep sequencing to examine the sequence dynamics and fitness changes of DENV populations adapting to human and mosquito cells, identifying the contributions of beneficial and deleterious mutations in shaping the fitness landscape driving host-specific paths of viral adaptation. We find DENV phenotypic adaptation is best described by the collective fitness contributions of all the alleles present in the population. Accordingly, while increased fitness during adaptation to each host is driven by host-specific beneficial mutations, it is reduced by the consistently replenished genetic load of deleterious mutations. Of note, host-specific beneficial mutations are in discrete regions across the genome, revealing molecular mechanisms of adaptation. Some of these clusters comprise phenotypically redundant mutations that may provide evolutionary robustness to transmission bottlenecks. Our results also suggest DENV adaptation is facilitated through variation in intrinsically disordered protein regions while transmembrane and structured domains evolve under stronger biophysical constraints. Importantly, the adaptation strategies uncovered in our simple system mirror macro-evolutionary changes observed across DENV serotypes and Zika virus and may suggest general principles of evolvability in arbovirus evolution.

## INTRODUCTION

The great evolutionary capacity of RNA viruses, driven by high mutation rates, allows them to adapt to their hosts and overcome barriers to infection ^2–5^. Understanding the molecular mechanisms of viral adaptation can reveal central aspects of host tropism and viral pathogenesis, as well as uncover fundamental principles governing molecular evolution. Arboviruses such as dengue (DENV), Zika (ZIKV), and chikungunya (CHIKV), which alternate between vertebrate and insect hosts, are a significant cause of disease globally. With half of the world’s population exposed, DENV alone causes approximately 100 million infections and 10,000 deaths annually ^1^.

Characterizing the genetic and evolutionary mechanisms underlying arboviral adaptation to the distinct environments of each host is key to understanding their emergence and spread. The vertebrate and invertebrate hosts of arboviruses differ significantly in temperature, cellular environment, and modes of antiviral immunity, which shape the evolutionary landscapes of viruses. Several studies are beginning to reveal how arboviruses relate to these alternative landscapes, identifying mutations that confer increased fitness in each host^6–11^. However, we still lack a comprehensive picture of the alternative fitness landscapes of any arbovirus defined by the human and insect host environments.

RNA viruses exist as a dynamic population of co-circulating mutant genotypes surrounding a master sequence^12^. While the distribution and dynamics of minor alleles are thought to play important roles in population fitness, adaptation and disease ^13–19^, technical limitations of sequencing resolution have restricted the analysis of experimentally evolved virus populations to allele frequencies greater than 1 in 100 or 1 in 1000. Recent developments in high-accuracy sequencing approaches ^20,21^, such as Circular Sequencing (CirSeq), which can detect alleles as rare as 1 in 10^6^ in frequency, allow us to probe much deeper into the full spectrum of diversity in a viral population. The ability to trace the evolutionary dynamics of individual alleles from their genesis at the mutation rate to their eventual fate in a given experiment, allows the description of viral fitness landscapes with unprecedented detail.

Here, we use CirSeq to characterize DENV adaptation to human and mosquito cells. By tracing individual allele trajectories for almost all possible single nucleotide variants across the DENV genome, we estimated the influence of positive and negative selection in shaping the evolutionary paths of DENV in the distinct environments. Analysis of the allele repertoire contributing to viral population fitness reveals the cumulative role of low-frequency alleles during adaptation. Moreover, we find that adaptation relies on host-specific beneficial mutations clustered in specific regions of the DENV genome. These regions are enriched in structural flexibility and are also sites of variation across naturally occurring DENV and ZIKV strains. By uncovering genetic and biophysical principles of DENV adaptation to its two hosts, our analysis provides insights into flaviviral evolution and reveals parallels between long and short-term evolutionary scales.

## RESULTS

### Phenotypic characterization of DENV populations adapting to human or mosquito cells

Two simple models could describe how arboviruses cycle between their alternative host environments (Fig. 1a). First, the viral genome could have overlapping host-specific fitness landscapes; in this case transmission would not involve significant trade-offs. Alternatively, the virus may have distinct host-specific landscapes with offset fitness maxima. To characterize the relative topography of the adaptive landscapes of DENV in vertebrate and invertebrate hosts (Fig. 1a), we performed an evolution experiment (Fig. 1b). Clonal, plaque-purified dengue virus (DENV serotype 2/16681/Thailand/1984) was serially passaged in the human hepatoma-derived cell line Huh7 or the *Aedes albopictus*-derived cell line C6/36, for nine passages. To maintain a constant, effective population size and to control the influence of drift due to population bottlenecks, a viral inoculum of 5×10^5^ focus forming units (FFU) from the previous passage was used to initiate the next passage by infecting 6×10^5^ cells at an MOI of 0.1. To distinguish host-specific versus replicate-specific events, we carried out two independent and parallel passaging experiments in each cell line (Series A and B, Fig. 1b).

**Fig. 1.**
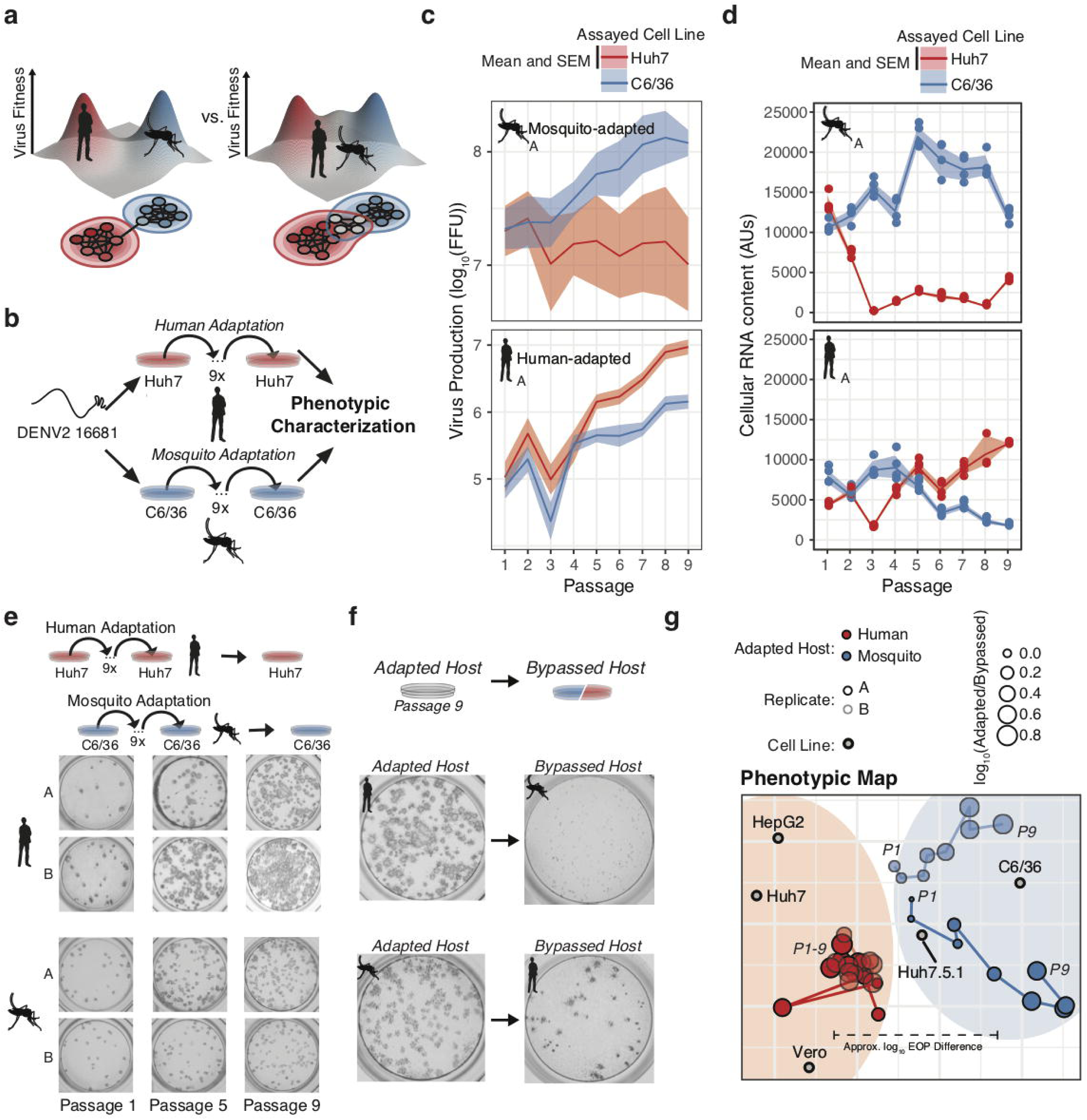
Dengue navigates distinct fitness landscapes in its alternative hosts. (a) Two potential models of the genotype-fitness landscape and mutational network in alternative arboviral hosts. The relative topography of the viral genotype-fitness landscape determines the extent of evolutionary trade-offs associated with transmission, and the paths through the mutational network toward host adaptation, and the proportion of genotypes viable in the alternative host environment (grey nodes). (b) Outline of our *in vitro* DENV evolution experiment. Dengue virus RNA (Serotype 2/16881/Thailand/1985) was electroporated into mosquito (C6/36) or human cell lines (Huh7), and the resulting viral stocks were passaged at fixed population size (MOI = 0.1, 5E5 FFU/passage) for nine passages in biological duplicates. After passage, viruses were characterized for phenotypic measures of adaptation. (c) Viral production assays comparing mosquito-adapted (top panel) and human-adapted (bottom panel) DENV populations. Adapted populations show increased virus production on their adapted hosts. Biological replicate A is shown for all experiments. (d) Analysis of viral RNA content by qRT-PCR. Cellular DENV RNA is significantly decreased when adapted lines are propagated on the by-passed, alternative host. Lines and shading represent the mean and standard deviation of 4 technical replicates, respectively. Biological replicate A is shown for all experiments. (e) Focus forming assays of the adapted populations over passage. Focus size increased markedly throughout passage on the adapted host. (f) Focus assays of the P9 virus on the adapted (left) and by-passed (right) host. Changes in focus size and morphology suggest evolutionary trade-offs between the alternative hosts. (g) *Tropic cartography* of DENV *in vitro* host adaptation. A two-dimensional embedding of the relative titer (mean of 4 replicates), or efficiency of plating (EOP), of the adapted populations (red and blue trajectories) relative to 5 assayed cell lines (grey points). Movement corresponds to a change in EOP over passage on a log scale. The size of the red and blue points indicate the ratio of the populations’ titer on the adapted and bypassed cell lines to illustrate the change in EOP over passage on the adapted lines.

The fitness gains associated with adaptation were assessed phenotypically by measurements of virus titer (Fig. 1c and Extended Data Fig. 1a), intracellular vRNA content (Fig. 1d and Extended Data Fig. 1b), and focus size and morphology (Fig. 1e and Extended Data Fig. 1c and d) for each viral population in the passaged host cell. All of these fitness measures increased over time for the passaged host, indicating significant adaptive evolution throughout the experiment. We quantified fitness trade-offs in parallel by carrying out the same measurements in the alternative (by-passed) host cell. In agreement with previous studies ^7,22–27^, passaging on one host cell line was accompanied by a concurrent loss of fitness in the alternative host cell line (Fig. 1c and Extended Data Fig. 1e). For instance, the human-adapted virus showed a uniform small focus phenotype when plated on mosquito cells (Fig. 1f). In contrast, mosquito-adapted populations formed fewer foci in human cells (Fig. 1f). Mosquito-adapted populations exhibited a heterogeneous focus phenotype, with small and large foci, suggesting they contain distinct variants differentially affecting replication in human cells.

We further assessed the evolutionary trade-offs during host adaptation by comparing the relative titers of all the passaged populations in both the original and the alternative host cells, Huh7 and C6/36. We also measured viral titers on human Huh7.5.1 cells, an Huh7-derived line lacking the RIG-I antiviral signaling pathway, as well as human hepatoma-derived HepG2 cells, and Africa Green Monkey epithelial-derived Vero cells (Fig. 1g and Extended Data Fig. 1d). For each passage, viral titers were normalized to that obtained in the adapted (original) host cell line, to yield the efficiency of plating (EOP) (individual EOP plots shown in Extended Data Fig. 1d). The large number of comparisons in 2D space were visualized using an embedding technique that summarizes the relative EOP as an approximate distance in two dimensions. This approach is similar to the technique of antigenic cartography used to describe antigenic evolution from pairwise measurements of antibody neutralizing titers ^28^. The movement of the sequenced viral populations (Fig. 1g, red and blue lines) relative to the placement of the cell lines (grey circles) reflects the change in relative titer between both cell lines. Human-adapted viruses exhibited similar titers in human- and primate-derived cell lines, resulting in their clustering separately from the titers in mosquito-derived C6/36 cells. The movement of C6/36-adapted populations (blue line) towards the C6/36 cell line reflects the significant mosquito-specific adaptation in these populations. Notably, RIG-I-deficient Huh7.5.1 cells placed intermediate to the other primate and insect cell lines, emphasizing the role played by innate immune signaling in the trade-offs associated with host adaptation.

### Characterizing genotypic changes in adapting DENV populations

To determine the genotypic changes associated with host cell adaptation, we subjected all viral populations to CirSeq RNA sequencing^21,29^, With an error rate of less than 1 in 10^6^, CirSeq yielded an average of approximately 2×10^5^-2×10^6^ reads per base across the genome for each viral population in our experiments (Fig. 2a)^20,29^. This depth permits the detection of alleles as rare as 1 in 60,000-600,000 genomes (Fig. 2b).

**Fig. 2.**
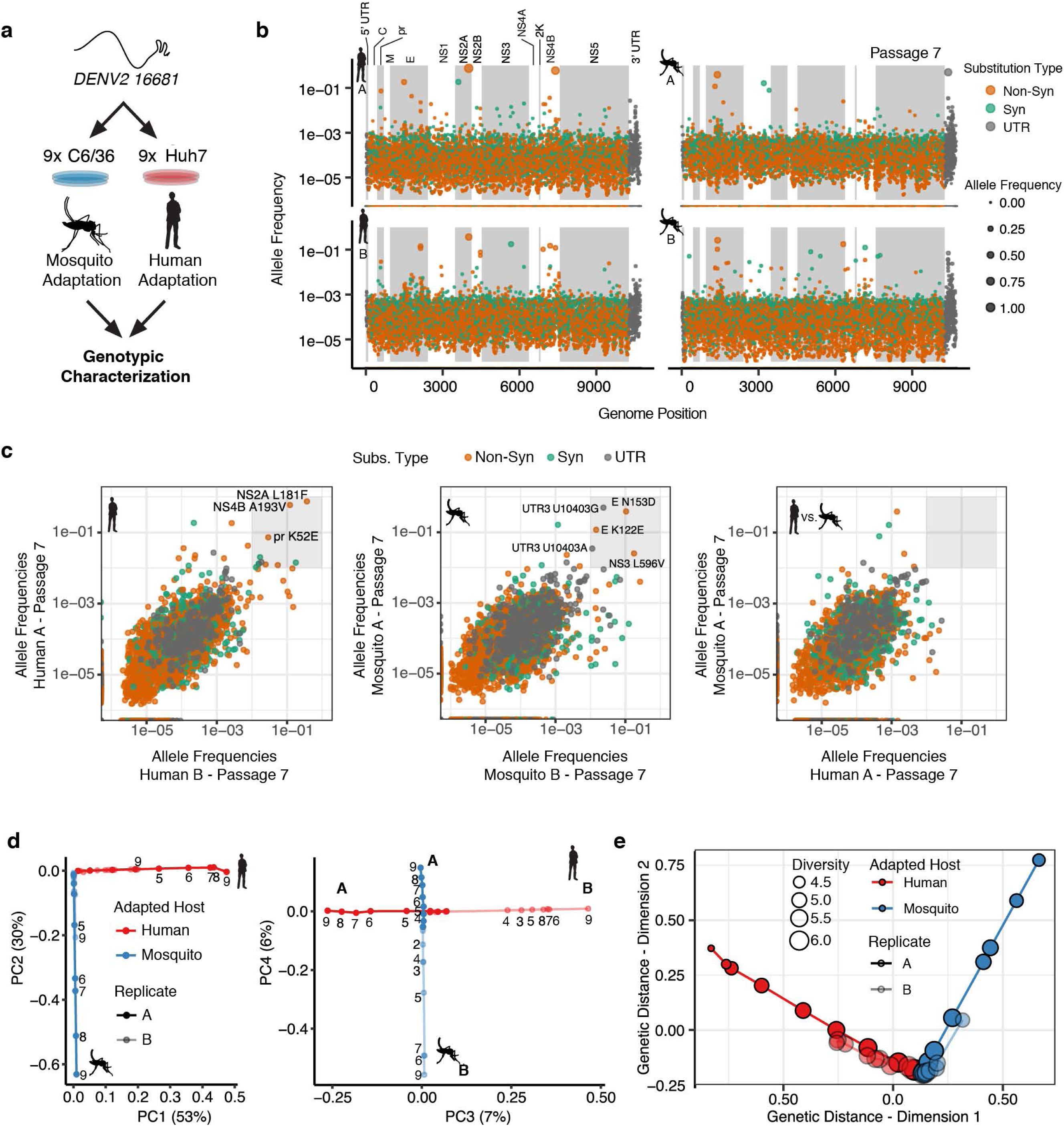
Adapting viral lineages show host-specific patterns of genetic variance. (a) Adapted viral populations were subject to genotypic characterization by ultra-deep sequencing using the CirSeq procedure. (b) Plots of allele frequency across the viral genomes for all four viral populations at passage 7. Alleles are colored by mutation type (Nonsynonymous, Orange; Synonymous, Green; Mutations in the untranslated region (UTR), Dark Grey). Shaded regions denote mature peptide boundaries in viral ORF. (c) Scatter plots comparing allele frequencies between adapted populations of human- and mosquito-adapted dengue virus. Replicate host-adapted populations share multiple high-frequency non-synonymous mutations, but populations from alternative hosts do not (grey square, >10%). (d) Dimension reduction of the allele frequencies by principal components analysis summarizes the host-specific patterns of variance (left), and the replicate-specific differences in genetic variability over passage (right). (e) A two-dimensional embedding of the pairwise genetic distances between the sequenced viral populations (Weir-Reynolds Distance) by multidimensional scaling. The viral populations (red and blue trajectories) project out from the founding genotype in orthogonal and host-specific directions. The size of the points corresponds to the diversity (computed as Shannon’s Entropy).

We next compared the mutational spectra in the sequenced populations (Passage 7 shown in Fig. 2b. No mutations reached fixation in the adapting populations, but many alleles rapidly increased in frequency with passage number (Extended Data Mov. 1). Comparison of the allele frequencies in the two independent passage series A and B revealed that the replicate populations (Fig 2c *i* or *ii*) shared more high-frequency mutations (defined here as 1% allele frequency) than populations passaged in different hosts, which shared no high frequency mutations (Fig. 2c *iii*). Human cell-adapted lineages shared high-frequency mutations in NS2A and NS4B, while mosquito replicates shared high-frequency mutations in E, NS3, and the 3′ UTR.

To better visualize the high-dimensional temporal dynamics of adaptation (Extended Data Mov. 1), we employed two alternative dimension reduction approaches that summarized the population sequencing data. Principal components analysis (PCA) quantifies the patterns of allele frequency variance between the populations, identifying independent patterns of variance (Fig. 2d and Extended Data Fig. 2a and b). Multidimensional scaling (MDS) examines the genetic distance and divergence of the populations over time (Fig. 2e). Both analyses demonstrate that the serially passaged DENV populations exhibit host-specific patterns of variation, revealing host-specific changes in genotypic space over passage in response to selection (Fig. 2d and e).

The PCA revealed the contribution of host-specific and replicate-specific changes in the viral population. The first four components of the PCA explained 96% of the observed allele frequency variance in the experimental populations. The first two components, which explain 83% of the observed variance (Fig 2d *i* and Extended Data Fig. 2a), partitioned the viral lineages along two orthogonal, host-specific paths, radiating outward in order of passage number from the original WT genotype (Fig. 2d *i*). The third and fourth components in the PCA explained 13.4% of the observed variance and further partition each lineage along orthogonal, replicate-specific axes (Fig. 2d *ii* for human series A and B and Fig. 2d *iii* for mosquito series A and B). The PCA-derived scores for individual alleles in component space summarized their contribution to the host- and replicate-specific dynamics (Extended Data Fig. 2b). The 3′ UTR and E contained the strongest signatures of mosquito-specific adaptation. Human-specific alleles were distributed across the genome, including nonsynonymous substitutions in E, NS2A, and NS4B (Extended Data Fig. 2b). Replicate-specific PCA components also highlighted clusters of alternative alleles in E and the 3’ UTR.

Multidimensional scaling (MDS) allows the embedding of the multidimensional pairwise distances between populations into two dimensions, providing a complementary view of the genetic divergence of the populations. Like PCA, this visualization also captures the orthogonal host-specific adaptive paths traveled by the populations (Fig 2e). In general, passage also corresponded to a loss of diversity in the population, measured as Shannon’s entropy (Fig. 2e).

The finding that reproducible population structures emerge during DENV adaptation to each host resonates with theoretical predictions that large viral populations will develop equilibrium states, commonly referred to as quasispecies, deterministically based on the selective environment ^30,31^. The host-specific composition of these populations likely reflects the differences in the selective environments that determine host range and specificity, prompting us to dissect their composition further.

### Fitness landscapes of DENV adaptation to human and mosquito cells

The concept of fitness links the frequency dynamics of individual alleles in a population with their phenotypic outcome, i.e., beneficial, deleterious, lethal, or neutral. Lethal and deleterious alleles are held to low frequencies by negative selection, while beneficial mutations increase in frequency due to positive selection (Fig. 3a). Observing the frequency trajectory of a given allele over time relative to its mutation rate enables the estimation of its fitness effect.

**Fig. 3.**
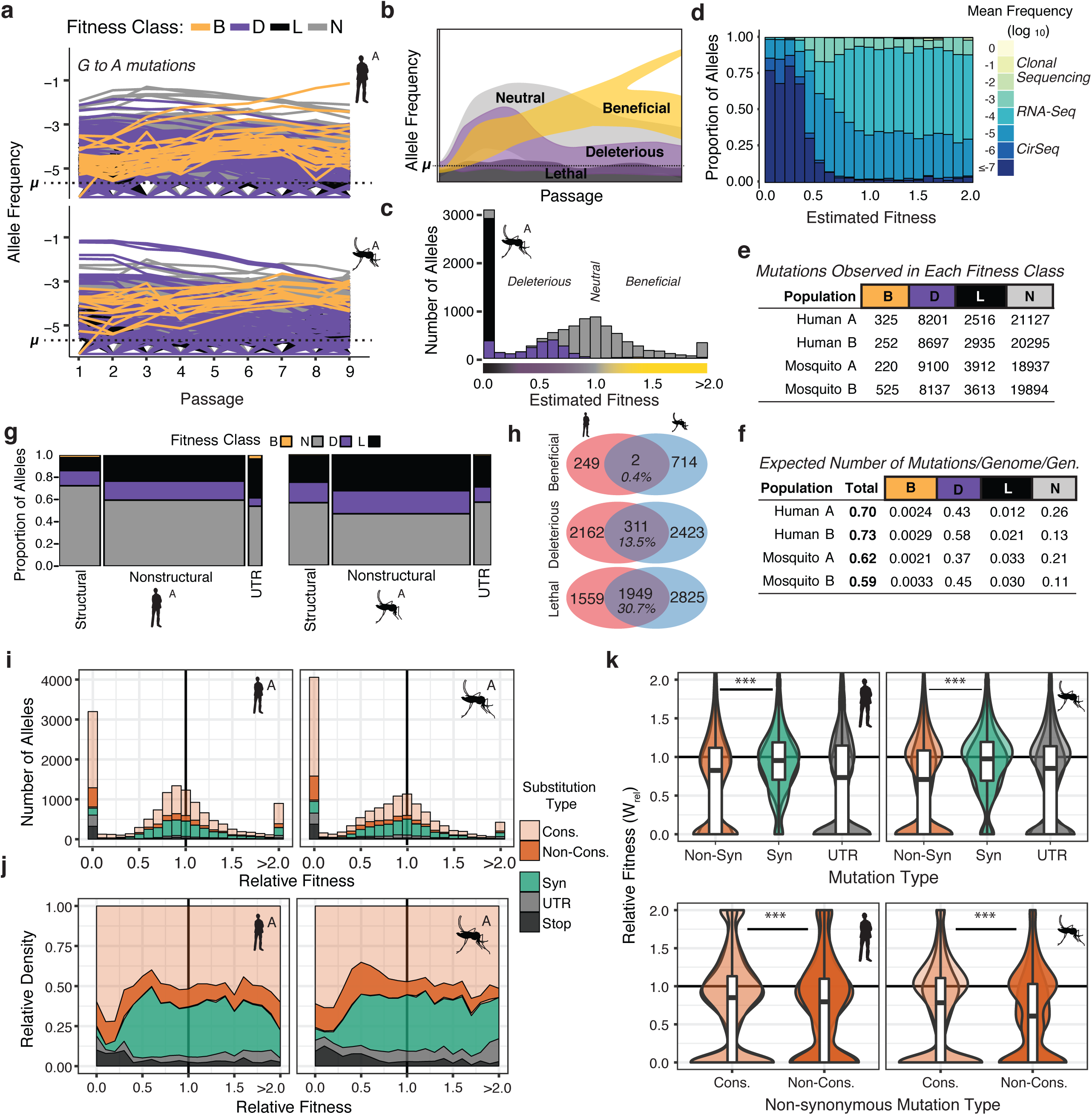
The distribution of fitness effects reveals patterns of evolutionary constraint. (a) The frequency trajectories for G to A mutations in the adapting populations determined by CirSeq. Colors represent the classification of each allele as beneficial, deleterious, lethal, or neutral (not statistically distinguishable from neutral behavior) (b) Schematic illustrating the expected frequency behavior of specific fitness classes relative to their corresponding mutation rate, *μ*. Changes in allele frequency between passages are used to estimate the fitness effects of individual alleles in the population (see Methods). (c) Histogram showing the distribution of mutational fitness effects (DMFE) of DENV passaged in mosquito cells. The data shown are from mosquito A and represent the high confidence set of alleles (see text). The fitness classifications of alleles in each bin, based on their 95% confidence intervals, is indicated by the fill color. (d) The relative density of each mutation type across the fitness spectrum illustrates the sequencing depth necessary to observe regions of the fitness spectrum. Fill color represents the average frequency of the mutation over passage. (e) Tabulation of all alleles by fitness class. (f) Estimate of the genomic mutation rate per genome per generation (“Total”), and fitness class-specific mutation rates (‘B’, ‘D’, ‘N’, and ‘L’). (g) Area plot showing the fitness effects associated with mutations in structural, non-structural, and UTR regions of the DENV genome. The relative width of the columns indicates the number of alleles in each class, the relative height of the colored regions indicates the proportion of alleles of a given class. (h) Venn diagrams showing the number of mutations identified as beneficial, deleterious, or lethal in the high confidence set alleles (see methods). (i) Histograms of the distribution of mutational fitness effects (DMFE) broken down by mutation type. (j) Density plot of the relative density of mutation types across the DMFE to emphasize the local enrichment of specific classes. (k) Violin plots showing the relative fitness of nonsynonymous and synonymous mutations, and those in the viral UTRs. Nonsynonymous mutations can further be partitioned into conservative and non-conservative classes, which differ significantly in fitness effect.

We took advantage of the resolution of CirSeq to estimate the substitution-specific per-site mutation rates for DENV using a previously described maximum likelihood (ML) approach (Extended Data Fig. 3a)^20^. These estimates, ranging between 10^−5^ to 10^−6^ substitutions per nucleotide per replication (s/n/r) for each substitution, agreed well across populations. C-to-U mutations occurred at the highest frequency, approximately 5×10^−4^ s/n/r in all populations, potentially reflecting the action of cellular deaminases (e.g., APOBEC3 enzymes ^32^). The genomic mutation rate, substitutions per genome per replication (s/g/r) (⎧g), was calculated by taking the sum of the ML mutation rate estimates of all single-nucleotide mutations across the genome, yielding ⎧g estimates of 0.70 and 0.73 s/g/r for mosquito populations and 0.61 and 0.60 s/g/r for human populations, consistent with genomic mutation rate estimates for other positive-strand RNA viruses (Extended Data Table 3) ^33^.

Using a model derived from classical population genetics (Fig. 3b), we next generated point estimates and 95% confidence intervals of relative fitness (ŵ) for each possible allele in the DENV genome (Fig 3c and d). The distribution of mutational fitness effects (DMFE) is commonly used to describe the mutational robustness of a given genome (Fig. 3c and Extended Data Fig. 3b). Describing the full spectrum of the DMFE, including the large subset of deleterious alleles, requires significant sequencing depth to establish each allele’s behavior relative to its mutation rate (Fig. 3b). The vast majority of alleles cannot be detected by clonal sequencing or conventional deep-sequencing approaches (Fig. 3d and Extended Data Fig. 3c), which can only detect a few high frequency beneficial and neutral mutations. Due to its low error rate, CirSeq reveals a much richer fitness landscape of adapting DENV populations (Fig. 3d), enabling us to examine the role of lower frequency alleles, such as deleterious mutations evolving under negative selection, on virus evolution.

The DMFEs of DENV exhibited bimodal distributions with peaks at lethality (ŵ =0) and neutrality (ŵ =1.0), and a long tail of rare beneficial mutations (ŵ>1.0), similar to what is observed for other RNA viruses ^20^. The 95% CIs of these ŵ fitness estimates were used to classify individual alleles as beneficial (*B*), deleterious (*D*), lethal (*L*), or neutral (*N*) (Fig. 3e and Extended Data Fig. 3a). Alleles with fitness 95% CI maxima equal to 0 were classified as lethal alleles; these never accumulate above their mutation rate due to rapid removal by negative selection (Fig. 3b and c, black). Alleles with an upper CI higher than 0 but lower than 1.0 were considered deleterious (Fig. 3b and Extended Data Fig. 3b, purple). Alleles with a lower CI greater than 1.0 were classified as beneficial; these accumulate at a rate greater than their mutation rate due to positive selection (Fig. 3b and Extended Data Fig. 3b, yellow). Alleles whose trajectories could not be statistically distinguished from neutral behavior (ŵ =1.0) are referred to as ‘neutral’ (Fig. 3b and e, and Extended Data Fig. 3b, gray) ^20^.

The genomic mutation rate represents the rate at which novel mutations enter the population (Fig. 3f, “Total”). To understand the expected fitness of new mutations, we used fitness classifications for all 32,166 possible single nucleotide variant alleles (Fig 3e) to estimate the genomic beneficial, deleterious, and lethal mutation rates (Fig. 3f). These estimates indicate that the virus maintains a significant deleterious genetic load due to the high rate at which deleterious and lethal mutations flow into the population. A given DENV genome will accumulate a deleterious mutation in 40-50% of replications, compared to a 0.2-0.3% chance of accumulating a beneficial mutation in any replication (Fig 3f).

### Defining Constraints Shaping the DENV Fitness Landscape

We compared the proportion of mutations in each fitness class for structural and non-structural regions of the viral polyprotein. A high confidence set of 13-14,000 alleles in each population was chosen based on sequencing depth and quality of the fit in the ŵ fitness estimates across passages. There were striking differences in the distribution of lethal and deleterious mutations in distinct regions of the genome (Fig. 3g). Non-structural proteins were significantly enriched in deleterious and lethal mutations compared to structural proteins. The increased mutational robustness of DENV structural proteins compared to non-structural ones contrasts with poliovirus, where structural proteins are less robust to mutation that nonstructural proteins ^20^. These differences may arise from the distinct folding and stability constraints of the enveloped and non-enveloped virion structures. We also find the viral UTRs exhibit host-specific patterns of constraint, consistent with their host-specific roles in the viral life cycle ^7,34,35^. In human cells, the DENV UTRs were more brittle but also contained more beneficial alleles than in mosquito adapted populations, suggesting strong selection.

In contrast to beneficial mutations, which were largely host and replicate specific, deleterious and lethal mutations exhibited significant overlap between the two hosts (Fig. 3h and Extended Data Fig. 3d), indicating common constraints on viral protein and RNA structures and functions in the two host environments. These biophysical constraints were further examined by evaluating how specific mutation types contribute to the viral fitness landscape (Fig 3i-k). As expected, synonymous mutations tended to be more neutral than non-synonymous mutations, which exhibited a bimodal distribution of fitness effects (Fig 3k). We further partitioned non-synonymous mutations into conservative substitutions (Fig. 3i-k, “Cons.”), which do not significantly change the chemical and structural properties of sidechains, and non-conservative which do (Fig. 3i-k, “Non-cons.”)^36^. Non-conservative changes exhibited a significantly greater deleterious fitness effects that conservative changes, emphasizing the impact of biophysical properties on fitness effects as well as the sensitivity of our approach to uncover these differences ^36^. As expected, lethal alleles were enriched in nonsense mutations (Fig. 3j, “Stop”) as well as nonsynonymous substitutions (Fig. 3j). These findings reveal the structural biophysical constraints shaping the DENV adaptive landscape and constraining viral diversity.

### Linking population sequence composition to experimental phenotypes

Examining the allele fitness distribution in DENV populations as a function of passaging in human or mosquito cells revealed a ‘fitness wave’ occurring over the course of the adaptation experiment (Fig 4a; Extended Data Fig 4a). In early passages, the population is dominated by neutral and deleterious mutations that arise during the initial expansion. In later passages, when rare beneficial mutations begin to accumulate under positive selection, we observe a concurrent loss of deleterious and neutral mutations, likely driven out by negative selection. However, because most mutations arising during replication are deleterious or neutral (Fig. 3f), their genetic load is never fully purged from the viral populations.

**Fig 4.**
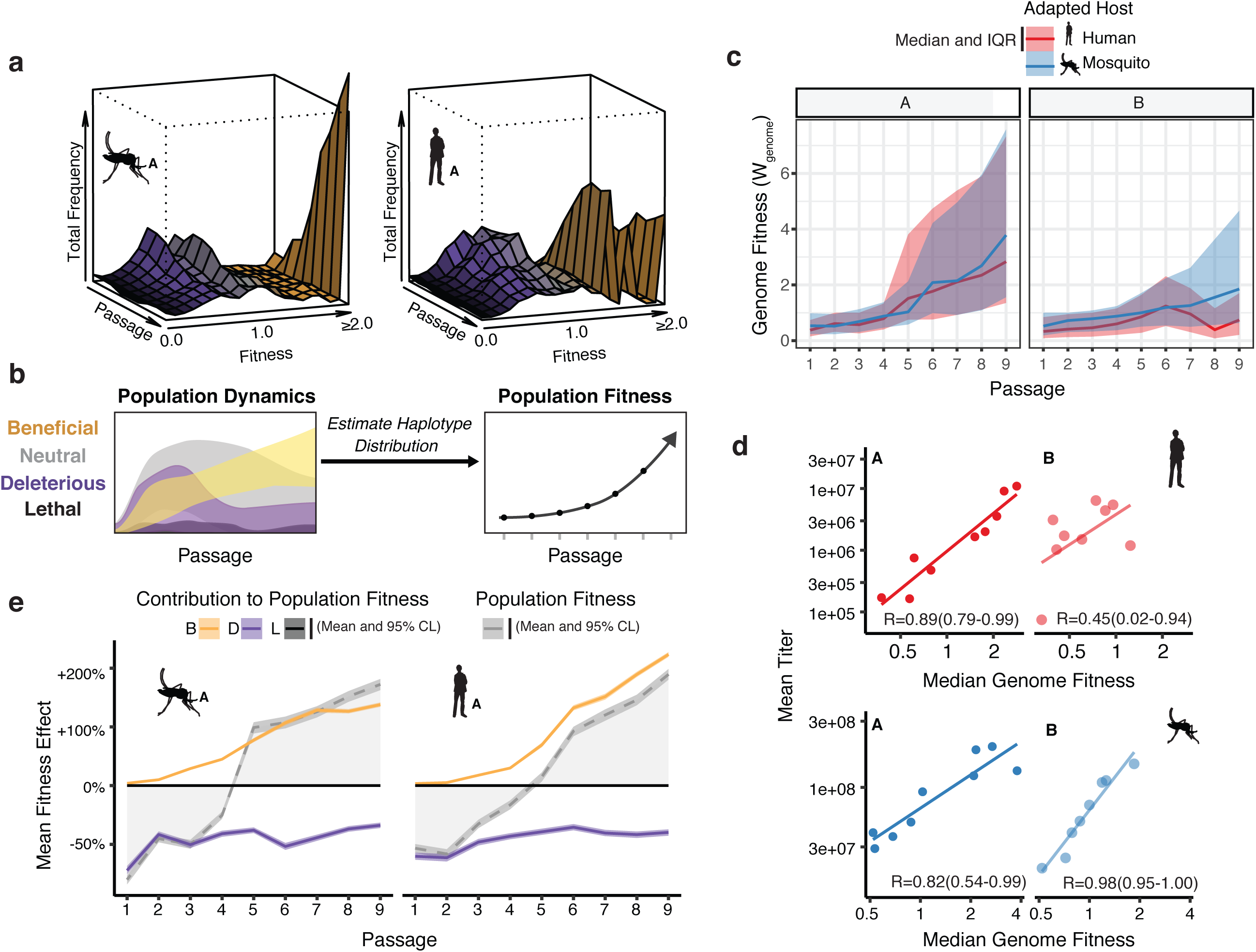
Connecting global evolutionary dynamics and population fitness. (a) The surface plot of the ‘fitness wave’ illustrates the change in frequency of alleles in each fitness bin throughout a passage experiment. The height of the surface represents the sum of frequencies of alleles in a given bin. Deleterious and neutral mutations (purple and grey regions) make up a large proportion of the population early in the experiment. They are mostly, but not entirely, driven out by beneficial mutations (yellow) in later passages. (b) The aggregate effect of the fitness wave is an increase in the mean population fitness, a weighted average of the fitness of individual genomes in the population. To estimate the distribution of genome fitness in the sequenced population in the absence of haplotype information, we estimated a collection of plausible haplotypes by sampling from the empirical frequencies of individual mutations and calculated the fitness of each assembled genome as the product of ŵ estimates across all positions (c) The median and interquartile range of genome fitness (*W*) of 50,000 simulated genomes sampled from the empirical allele frequencies determined from each sequenced population. (d) Correlation plots comparing the median genome fitness (*W*) of the sampled populations and mean virus titer correspond well to each other. (e) Line plots showing the influence of beneficial (yellow), deleterious (purple), and lethal (black) mutations on the mean fitness of the population (grey line). The shaded area represents the 95% confidence interval of the mean from 50,000 simulated genomes (see Materials and Methods).

The Fundamental Theorem of Natural Selection dictates that the mean relative fitness of a population should increase during adaptation ^37,38^. Given the low probability of acquiring multiple mutations per genome per replication cycle (Fig. 3f), individual alleles were treated as independent of each other in our previous analysis of fitness. However, linking the fitness of individual alleles to the dynamics of genomes, and estimating the aggregate effect of individual mutations, requires the estimation of haplotypes (*W*). To this end, we used the fitness values and frequency trajectories of individual alleles to estimate the aggregate changes in fitness in the populations over the course of adaptation to either host cell (Fig. 4b). For each population, we generated a collection of haplotypes, and corresponding genomic fitness values (*W*) that reflect the empirical allele frequencies determined by CirSeq (Fig 3). This simulated population was then used to explore the changes in distribution of *W* at the population-level over the course of passage. As expected, the median *W* of these simulated populations increased throughout passaging in a given cell type (Fig. 4c), consistent with the dynamics of the individual constituent alleles (Fig. 4a).

Next, we compared the genotype-based median fitness of the population (Fig. 4d) with the experimental phenotype observed in the corresponding viral population (Fig. 1). We chose mean absolute viral titers (Fig. 1c) as a gross measure of population replication fitness and adaptation to the host cell. We observed a striking correlation between viral titers and the calculated median *W* of each population, based on allele frequency trajectories alone (Fig 4d; R values ranging from 0.45 to 0.98). This correlation suggests that the comprehensive measurement of allele frequencies can capture the phenotype and adaptation dynamics of these virus populations.

The findings above indicate that a sequencing-based estimate of population fitness that incorporates the entire complement of alleles reflects experimental phenotypes in that population. We next examined the contribution of beneficial, lethal and deleterious mutations to mean population fitness. To this end, we again calculated the distribution of genome fitness for each adaptation lineage as described above, but taking into account the influence of only beneficial, deleterious or lethal alleles. Next, we compared these class-specific trajectories to the overall mean population fitness (Fig. 4e, broken grey line) in order to estimate the contribution of each class of mutation to the adaptation of the populations over passage (Fig. 4e). Beneficial mutations, although occurring relatively rarely, rapidly accumulate and drive the increase in the mean relative fitness (Fig. 4e, yellow line). In contrast, deleterious alleles, which individually are present at low frequencies but occur on 40-50% of genomes, contribute a significant mutational load across all passages (Fig. 4e, purple line). Accordingly, the burden of deleterious mutations reduces mean fitness by approximately 50% across all passages (Fig. 4e, purple line). Of note, lethal mutations (Fig. 4e, black line) exert minimal effect on the population because they are rapidly purged and remain very rare. These analyses reveal that both beneficial and deleterious mutations play important roles in the phenotypes of large, diverse viral populations. Importantly, they indicate that the high levels of deleterious mutations in a viral population exerts a persistent burden during adaptation. Sequencing approaches like CirSeq will be essential for comparing the mutational burden on different viral species to understand the balance between mutational tolerance and constraint on viral genomes.

### Molecular and structural determinants of dengue host adaptation

Analysis of the regions of the viral genome under positive and negative selection in each host provided insights into the molecular determinants of DENV adaptation. We calculated the mean fitness effect of non-synonymous and noncoding mutations in 21 nucleotide windows and mapped them onto the genome (Fig. 5a). Regions of evolutionary constraint, denoted by deleterious (purple) and lethal (black) mean fitness effects, were found throughout the genome, distributed similarly between the two hosts. These likely reflect general constraints on protein structure and function. For instance, regions in non-structural proteins NS3, NS4B, NS5, and in the UTRs shared regions of strong negative selection in both hosts, which may denote key structural and functional elements. In contrast, the patterns of positive selection along the genome were different between the two hosts (yellow points in Fig. 5a). Notably, many beneficial mutations were clustered at a few specific locations in the genome (yellow points in Fig. 5a), suggesting adaptation relies on hotspots of host-specific selection.

**Fig. 5.**
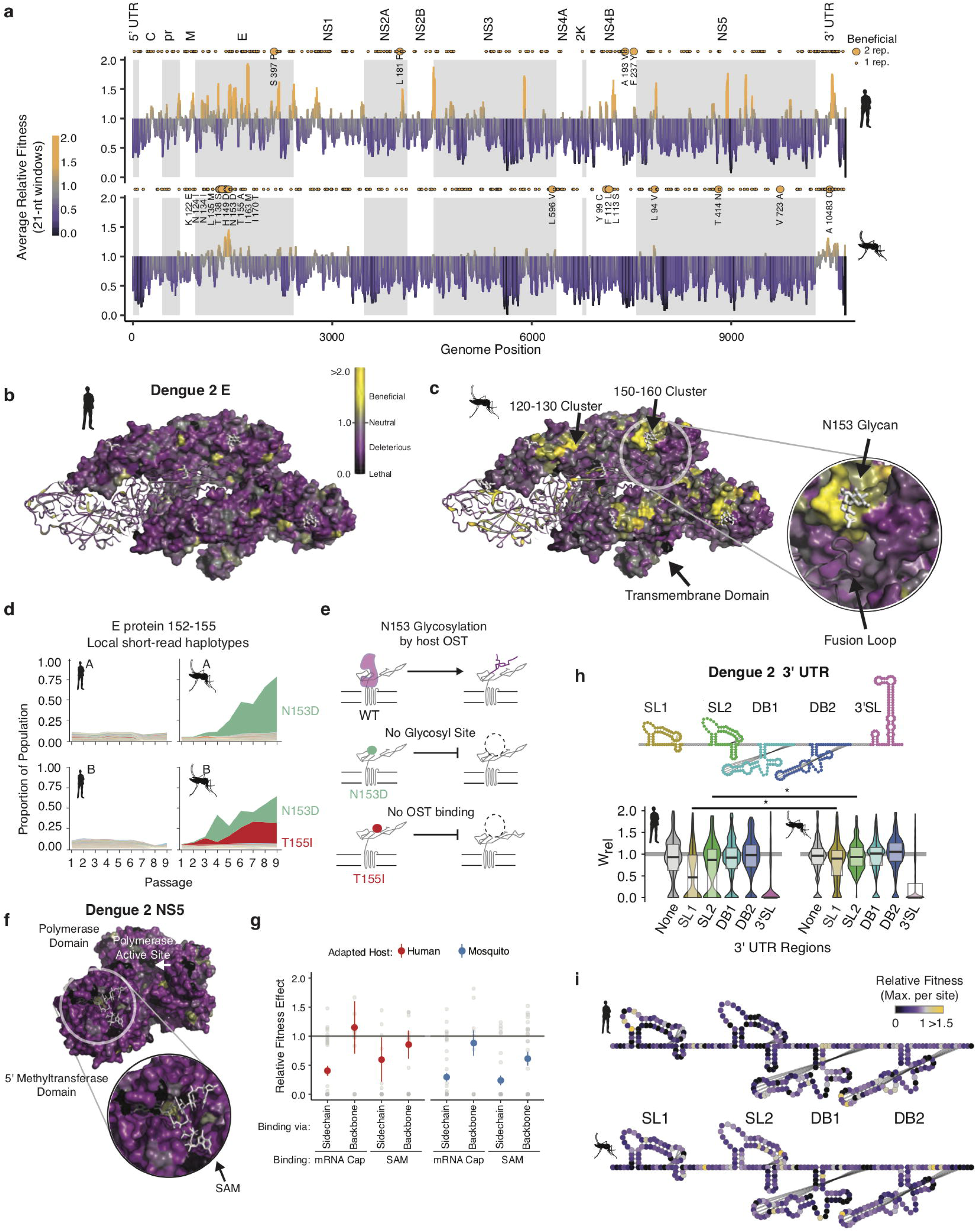
Structural analysis reveals hotspots of viral adaptation. (a) Barplot of the mean fitness effect of alleles in 21 nt windows across the DENV genome. Synonymous alleles are removed to emphasize the fitness effects of coding changes. Yellow points above each line denote the locations of beneficial mutations. Larger, labeled yellow points denote beneficial mutations identified in both replicates of host cell passage. (b) The empirical fitness estimates displayed on a trimer of envelope proteins in an antiparallel arrangement, similar to that found on the mature virion. To emphasize rare sites of positive selection, the color of each residue represents the maximum of the lower confidence intervals of fitness effect estimates at that site. Human-adapted populations show significant negative selection on the envelope protein surface, with no residues showing significant positive selection (yellow color). (c) Mosquito-adapted DENV exhibit two patches of pronounced positive selection on the exterior face of the virion (labeled 120-130, and 150-160). These clusters are absent from the human-adapted populations. Cluster *150-160* (Zoom), consists of a loop containing a glycosylation site at N153. This loop and glycan enclose the viral fusion loop of the anti-parallel monomer prior to activation and rearrangement in the endosome after entry. (d) Plots of the frequency of local read haplotypes for the area overlapping N153 and T155. Mutations at N153 (N153D) and T155 (T155I) are positively selected in mosquitoes, but never occur together on individual reads. (e) Schematic describing the phenotypically equivalent effects of the N153D and T155I mutations. These mutations block recognition and modification by the host oligosaccharyltransferase (OST), which initiates glycosylation. (f) CirSeq also reveals patterns of negative selection. Patches of significant evolutionary constraint can be seen around the methyltransferase active site highlighted by numerous positions with lethal fitness effects (Zoom). (g) Comparison of fitness effects of non-synonymous mutations targeting residues in NS5-MT that interact with the 5 (h) Insights into host-specific RNA structural constraints. Violin plot comparing the fitness effects of mutations in the stem-loop (SL) and dumb-bell (DB) structures of the 3′ UTR RNA of DENV2 shown in the schematic. Fitness effects of mutations in the conserved structures reveal differences in fitness effects associated with SLI and SLII in human- and mosquito-adapted dengue virus populations. (i) Nucleotide-resolution map of fitness effects on the viral 3′ UTR reveals regions of SL1 and 2 that are under tighter constraints in human passage.

To further analyze these “adaptation hotspots,” we mapped the allele fitness values on the three-dimensional structure of dengue protein E, which forms the outermost layer of the viral envelope (Fig. 5b and c). The clusters of adaptive mutations identified in mosquito cells were under negative selection in human-adapted populations (Fig. 5b). One mosquito-adapted cluster mapped to residues 150-160, surrounding the glycosylation site at N153 (Fig. 5c, zoomed region), which has been associated with mosquito adaptation ^39^. Closer examination of this loop (E152-155) revealed that different mosquito alleles created alternative substitutions, N153D and T155I, with identical phenotypic consequences, namely to abrogate N153 glycosylation. N153D eliminates the asparagine that becomes glycosylated, while T155I disrupts the binding of the oligosaccharyltransferase mediating glycosylation (Fig. 5d). Since both positively selected mosquito alleles disrupt NxT glycosylation at this site ^40^, it appears that eliminating this glycan moiety is beneficial in mosquitoes but not in human cells (Fig. 5d). Interestingly, glycosylation pathways diverge significantly between humans and insects, yielding very different final glycan structures ^41^. Since this loop is a primary site of structural variation in E proteins of dengue and related flaviviruses, including Zika virus ^42^, its diversification may reflect past cycles of host-specific selection acting on this region of E.

The ability of CirSeq to detect alleles at frequencies close to the mutation rate permits detection and quantification of negative selection, revealing sites that are critical for viral replication. Therapies targeting these highly constrained regions under strong negative selection may be less susceptible to resistance mutations. For instance, both the RNA polymerase and the methyltransferase active-sites of NS5 are enriched in lethal mutations in residues contacting the enzyme substrates (Fig. 5e). Further analysis of mutations in the methyltransferase residues contacting its ligands SAM and mRNA cap illustrates the power of our analysis. We find residues that contact the ligands through sidechain interactions are under strong negative selection, with most mutations highly deleterious. In contrast, residues that interact with the ligands through backbone interactions were relaxed in their fitness effects (Fig. 5f). Thus, such high-resolution evolutionary analyses could complement structure-based anti-viral drug design by identifying regions of reduced evolutionary flexibility, which may be less prone to mutate to produce resistance.

Our analyses also captured key differences in the evolutionary constraints on the viral 3′ UTR (in Fig. 5h). We find that stem-loop II and the nearly identical stem-loop I in the 3′ UTR show significant shifts in mutational fitness effects between human and mosquito cells (Figs. 5h and i). These stem-loops are conserved across flaviviruses and form a “true RNA knot,” capable of resisting degradation by the exonuclease XRN1^43–45^. While previous findings indicated that in mosquito DENV acquired deletions in these loops ^7^ our findings indicate DENV also accumulates highly beneficial mutations that alter stem-loop folding in mosquito, which may result in the accumulation of distinct small RNAs upon XRN1 exonucleolytic activity. Our ability to identify subtle shifts in the DMFE can thus point to molecular mechanisms of selection and adaptation.

### Defining design principles of dengue virus adaptability

The surprising finding that adaptive mutations cluster in specific regions of the genome suggests that adaptation operates through discrete elements. We next examined the structural and functional properties of these elements to better understand the design principles of DENV evolution.

The Dengue polyprotein consists of soluble and transmembrane domains. We found that transmembrane domains were depleted of beneficial mutations and enriched in lethal mutations (Fig 6a). Thus, despite differences in lipid composition of human and insect membranes ^46,47^, the transmembrane regions of DENV disfavor change during host cell adaptation. For non-transmembrane DENV regions, we found striking differences between structured domains and intrinsically disordered regions (IDRs) (Fig 6b). Beneficial mutations were highly enriched in IDRs, but not in structured regions (Fig. 6b). In contrast, lethal mutations were enriched in ordered domains, while strongly depleted from IDRs, highlighting the evolutionary constraints imposed by maintaining folding and structure.

**Figure 6:**
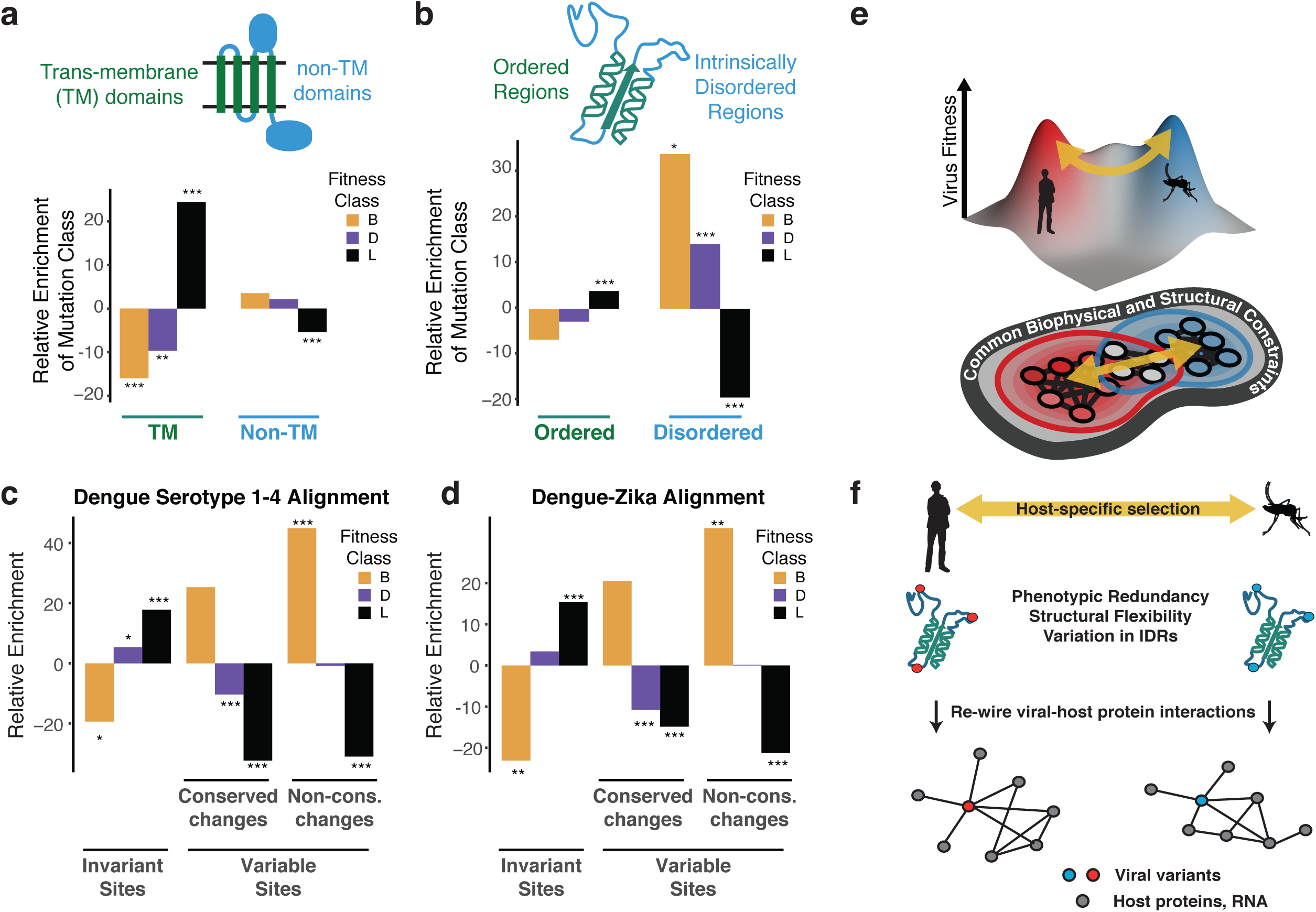
Biophysical and biological themes in DENV adaptation. (a, b) Distribution of mutations in regions with different biophysical characteristics. (a) The relative enrichment of each type of mutations (beneficial, deleterious, and lethal) in transmembrane (TM) regions versus non-TM regions (Fig. 2a), and disordered versus ordered regions (Fig. 2b). Relative enrichment is computed as the normalized difference in occurrence of this type of mutation in the specific region tested, and its occurrence across the entire polyprotein. Significance values (FDR-corrected, Fisher exact values) are shown (* = p<0.05, ** = p<0.01, *** = p<0.001). (c) Cross-dengue conservation. Distribution of mutations in regions with different levels of conservation across dengue virus strains. The relative enrichment of each type of mutations (beneficial, deleterious and lethal) in residues that are identical (“Invariant”), similar (“Conserved”) or dissimilar (“Variable”) across the four dengue strains. Relative enrichment was calculated s computed as the normalized difference in occurrence of this type of mutation in the specific region tested, and its occurrence across the entire polyprotein. Significance values (FDR-corrected, Fisher exact values) are shown (* = p<0.05, ** = p<0.01, *** = p<0.001). (d) Zika-Dengue conservation. Distribution of mutations in regions with different levels of conservation between DENV and ZIKV virus. The relative enrichment of each type of mutations (beneficial, deleterious, and lethal) in residues that are identical, similar, or dissimilar between the two viruses. Relative enrichment was calculated as the normalized difference in occurrence fraction of this type of mutation in this specific region and its occurrence across the entire polyprotein. Significance values (FDR-corrected, Fisher exact values) are shown (* = p<0.05, ** = p<0.01, *** = p<0.001). (e) Visualization of a simplified landscape of dengue host adaptation. The landscape is shaped by common biophysical and functional constraints that operate similarly in both hosts, defining the outline of the fitness landscape. Positive selection of host-specific phenotypes drives host adaptation. (f) Host adaptation is associated with trade-offs that form a bottleneck to transmission. This bottleneck is relaxed by phenotypic redundancy and structural flexibility at key hotspots of adaptation.

These analyses uncover underlying principles for how the DENV sequence space evolved to facilitate access to fitness peaks in its two distinct host cell environments. We find that a large fraction of the DENV genome sequence space does not respond to host-specific pressures. Instead, adaptation is mediated by changes in structurally flexible regions. Thus, we find that transmembrane and structured domains are not subject to optimization through host-specific beneficial mutations, indicating these regions reside at a trade-off point for efficient replication in both hosts. Interestingly, adaptation to each host cell operates primarily through variation in flexible, surface-exposed disordered regions. IDRs have few structural constraints and tend to mediate protein-protein and protein-RNA interactions, making them well suited for the evolutionary remodeling of virus-host-specific networks ^48,49^.

We next examined whether the patterns observed in our short-term reductionist adaptation study relate to patterns observed in long-term evolution (Fig. 6c). Sequence alignments of all four major DENV serotypes were used to classify amino acid residues that are invariant across the four strains, those with conservative substitutions that maintain chemical properties, and those with highly variable non-conservative substitutions. Strikingly, when compared to the fitness classes derived in our study, the occurrence of lethal, detrimental, and beneficial mutations mirrored the evolutionary conservation and variance across DENV serotypes (Fig. 6c). For instance, beneficial mutations in our study were strongly enriched in the regions of highest variation across DENV serotypes, while lethal mutations were enriched in residues that are invariant during evolution. These conclusions were supported when extending this analysis to include conservation between DENV and Zika virus (Figs 6d). Displaying agreement between the two compared evolutionary scales, we find that regions displaying higher constraints in long-term evolution are depleted of beneficial mutations and enriched in lethal mutations arising from our short-term cell culture analysis. In contrast, regions of higher variation between viral species have fewer lethal mutations and a higher level of beneficial mutations. Thus, our simple cell culture paradigm uncovers patterns of conservation and adaptation that reflect evolvability design principles of flaviviruses cycling between human and mosquito hosts.

## DISCUSSION

Here we explored the dynamics and global fitness changes in DENV populations adapting to human and mosquito cells. Using high-resolution sequencing, we identify the contribution of beneficial and deleterious mutations shaping the evolutionary paths of DENV populations responding to host-cell specific selective pressures. Our analysis shows that DENV populations move in largely reproducible paths during adaptation to human or mosquito cells, driven by host-specific selection (Fig. 2). These host-specific population structures are defined by distinct genotype-fitness landscapes (Figs 1, 2), which collectively shift the population in sequence space and result in a concurrent increase in phenotypic fitness, as assessed by focus morphology and absolute titers (Fig. 1). Strikingly, simple models of mean population fitness derived from allele frequency measurements alone can predict these phenotypic changes (Fig. 4). This suggests the phenotype and evolutionary dynamics of a virus can be described by the fitness contributions of all alleles in the population. An insight of these analyses is that the large burden of detrimental mutations present in viral populations imposes a significant fitness cost. This observation resonates with the finding that the high mutation load of RNA viruses, while driving adaptation, places them at the “error-catastrophe threshold” ^3, 4, 14^. It will be interesting to understand how each type of mutation contributes to overall population fitness and in particular, how detrimental mutations reduce global replication fitness.

We find that adaptive mutations, including changes in coding and noncoding regions, are clustered in specific regions across the DENV genome. Examination of mosquito specific alterations in a glycosylation site in protein E and in the 3’ UTR suggest that mutations in these clusters lead to similar phenotypic outcomes. For instance, mutations that cluster in the 3′ UTR change the structure of stem-loop II, a site for gate-keeper mutations for mosquito transmission ^7,9^. Similarly, mutations that cluster in loop 150-160 in DENV E protein disrupt glycosylation in mosquito cells ^39^. This suggests that the process of host adaptation relies on the availability of multiple changes that can access a similar adaptive phenotype. We propose that the phenotypic redundancy of such mutations increases the mutational target size associated with key transitions necessary for adaptation, thereby partially relieving possible bottlenecks associated with transmission and early adaptation.

Our study highlights the crucial role of structural constraints in shaping DENV evolution. Adaptive mutations are largely excluded from transmembrane domains and structured regions in DENV proteins. Thus, structural integrity places significant constraints on variation within these regions. It is tempting to speculate that the sequence of these arboviral domains is poised at a compromise that optimizes function in the distinct environments of human and mosquito cells ^50–52^. Of note, our identification of highly constrained DENV regions, where most mutations are lethal, may uncover attractive targets for antivirals.

Beneficial mutations are enriched in flexible loops and intrinsically disordered regions of the DENV polyprotein (Fig 6b). The relaxed structural constraints of IDRs allow them to explore more mutational diversity without compromising protein folding or stability, thus enabling access to more extensive sets of adaptive mutations ^48,53^. Such plasticity may allow viral IDRs to rewire viral protein interactions with host factors, thereby driving adaptation to changing environments (Fig 6f).

Notably, the link between structural properties and fitness effects measured in our study mirrors sequence conservation and variation across natural isolates of DENV and ZIKV (Fig. 6c, d). This indicates that the relationships between adaptability, structural flexibility, and phenotypic redundancy uncovered here for DENV adaptation to cultured human and mosquito cells can inform on general principles in flavivirus evolution. While arbovirus that cycle between human and mosquito represent a more extreme case of host switching, most emerging viruses must adapt to changing environments during zoonotic transmission or intra-host spreading. We propose that our simple experimental approach can identify regions mediating evolutionary changes linked to diversification, tropism, and spread for a wide range of RNA viruses.

## METHODS

### Cells

Huh7, Huh7.5.1, HepG2, Vero cells were cultivated at 37°C and C6/36 cells at 32°C, respectively, as described previously ^54^.

### Viruses

DENV2 strain 16681 viral RNA generated by *in vitro* transcription was electroporated to produce progeny virus as described in ^54^. For passaging, DENV of 5 × 10^5^ FFU were serially propagated in one 10cm dish of Huh7 or C6/36 cells. The culture medium was collected before the cells showed a severe cytopathic effect (CPE). At each passage, virus titers in the supernatant were measured and adjusted for the next passage.

### Focus forming assay

Semi-confluent cells cultured in 48-well plates were infected with a limiting 10-fold dilution series of virus, and the cells overlaid with culture medium supplemented with 0.8% methylcellulose and 2% FBS. At 3 (Huh7) or 4 (C6/36) days post-infection, the cells were fixed by 4% paraformaldehyde-in-PBS, stained with anti-E antibody and visualized with a VECTASTAIN Elite ABC anti-mouse IgG kit with a VIP substrate (Vector Laboratories, Burlingame, CA USA). The entire wells of 48-well plates were photographed by Nikon DSLR camera D810, and each foci size was measured by image-J. Each experiment was performed in duplicate.

### Quantitative Real-Time PCR (qRT-PCR)

The intracellular RNAs were prepared by phenol-chloroform extraction. cDNA was synthesized from purified RNA using the High Capacity cDNA Reverse Transcription Kit (Life Technologies), and qRT-PCR analysis performed using gene-specific primers (iTaq™ Universal Supermixes or SYBR-Green, Bio-Rad) according to manufacturers’ protocols. Ct values were normalized to GAPDH mRNA in human cells or 18S rRNA in mosquito cells. qRT-PCR primers are listed in Table S1. Each experiment was performed in triplicate.

### CirSeq and Analysis of allele frequencies

For preparing CirSeq libraries, each passaged virus (6 × 10^6^ FFU) was further expanded in parental cells seeded in four 150 mm dishes. The culture medium was harvested before the appearance of severe CPE, and the cell debris was removed by centrifugation at 3,000 rpm for 5 min. The virion in the supernatant was spun down by ultracentrifugation at 27,000 r.p.m, 2 hours, 4°C and viral RNA was extracted by using Trizol reagent. Each 1µg RNA was subjected to CirSeq libraries preparation as described previously ^21^. Variant base-calls and allele frequencies were then determined using the CirSeq v2 package.

### Calculation of Relative Fitness

An experiment of *N* serial passages will produce, for any given single nucleotide variant (SNV) in the viral genome, a vector *X* of variant counts at each passage, *t*:

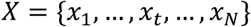

And, a vector Y containing the corresponding coverages at each passage, *t*:

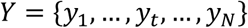

As explained previously^1^, the relative fitness of a SNV, *w*, at time *t* can be described by the linear model:

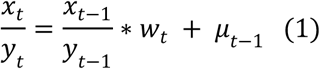

Where *μ*_*t*−1_ is the estimated mutation rate for the variant at time *t* − 1 (described previously^1^). This model requires only two consecutive passages to estimate a relative fitness parameter. However, to account for and quantify passage-to-passage noise in the estimates of relative fitness we used the values of *w* across passages to estimate the mean and variance of *w* for each SNV.

To account for genetic drift in our experiment, we used a similar approach as Acevedo et al., 2014. At each passage, a fixed number of focus forming units, *β*, are used to infect each subsequent culture. In each *β* virions, *b*_*t*−1_ of them will carry a given SNV. Therefore, 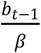 can be used to express the frequency of that SNV in the transmitted population1 which when substituted for the term, 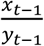, in the right side of equation (1) will yield:

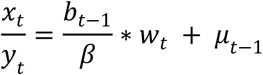

or:

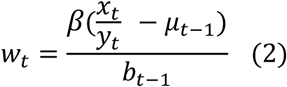

where:

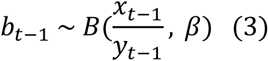

Given that *β* is constant (5×10^5^ FFU), we need only calculate *b*_*t*−1_ in order to estimate *w*_*t*_ values. Since we do not know the real value of *b*_*t*−1_ for any variant, especially for low frequency variants which are sensitive to bottlenecks, we need to estimate it. This can be done by sampling *m* times from equation (3). Such sampling is described by a Poisson distribution, then:

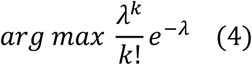

will give a maximum likelihood estimate for *λ* = *b*_*t*−1_, while the upper bound of *k* is given by *β*. Doing so for each *x* from time 1 to N-1 of gives a vector, 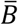, of *b* values: 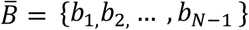. Finally we estimate N-1 *w* values by solving equation (2) using each element of 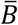. This gives a vector 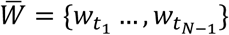.

Then, the slope of the linear regression over the cumulative sum of 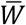 yields the estimated relative fitness, ŵ, of a given SNV. For this regression, we employed the Thiel-Sen regression method (implemented in the R package ‘deming’ ^55^), given that some of our vectors 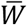 contains outliers as the result of 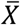 having zeros due to poor coverage. This regression will allow for the estimate to be robust to those outliers, to avoid classifying them as detrimental variants because spurious zeros. At the same time, for 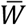 with a majority of zeros and some positive observations, that are likely to come from elements in *X* that are not significant (i.e. sequencing errors), the Thiel-Sen estimate will give more weight to the real zero values, classifying them as lethal or deleterious, and neglecting the effect of the positive elements in *W*. Finally, we also obtain an estimate of the 95% confidence interval by the procedure described previously ^56^ and implemented in the ‘deming’ package ^55^.

### Calculation of Mean Fitness

To estimate the effect of the observed population dynamics on the fitness of individual viral genomes and the distribution of genome fitness in the absence of haplotypic information, we generated a population of genomes sampled from the empirical allele frequencies. For each genome, sequence identity and corresponding fitness estimates, ŵ, at each position were randomly selected with a probability equal to its frequency in each sequenced population. Following sampling, the fitness of each genome in the sampled population was calculated as the product of the ŵ values across all positions:

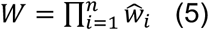

For all simulations, 50,000 genomes were sampled, and statistical analyses performed on the resulting populations; similar results were obtained with independent samples. To estimate the contribution of mutations in the individual fitness classes (i.e. beneficial, deleterious, or lethal) to the aggregate population fitness, sampling was performed as described above. Following sampling, the genome fitness, *W*, was calculated as described in equation (5), but only taking into account substitutions of a given fitness class.

#### Dimension reduction of genotypic data

Principal components analysis was performed on the unscaled population allele frequencies using the ‘princomp’ function in the R base ^57^. Calculation of Reynold’s Θ was performed using the adegenet ^58^ and poppR ^59^packages in R ^57^. Classical MDS (by Torgerson scaling ^60^) was performed to embed the pairwise Reynolds distances (Θ) ^61^ between the viral populations in 2-dimensions.

#### Dimension Reduction of Phenotypic Data

Stress Minimization by Majorization (implemented in *SMACOF* ^*62,63*^) was used for the ordination of cells and viruses based on empirical relative titer data. The input distance matrix was generated from the mean of log_10_ titer measurements for (N=4) focus formation assays on each passaged population on each of five cell lines: Huh7, Huh7.5.1, C6/36, HepG2, and Vero. Titer values were log_10_ transformed and subtracted from the maximum log_10_(titer) for each cell line to yield a matrix of Cell-to-Population distances, where the minimum distance represents the highest relative viability for each virus population.

### Structural Analysis

Fitness values for non-synonymous mutations were displayed on available dengue pdb structures using pyMol2 (Schrödinger). Data was aligned to structures using in-house scripts. Briefly, protein sequences for each chain in the PDB structure are mapped to the dengue 2 reference polyprotein sequence using the Smith-Waterman algorithm for pairwise alignment (implemented in ‘SeqinR’ package). To emphasize regions of positive selection, the values displayed on the structures represent the lower 95% confidence limit of the fitness estimate. Where multiple non-synonymous alleles could be mapped to a single residue, the maximum of the lower 95% confidence limits were displayed to emphasize the most significantly positively selected alleles at any position.

### Biophysical properties analyses

We have identified transmembrane regions using TMpred ^64^, taking regions with a score above 500 as *bona fide* transmembrane regions. For disorder prediction, we used IUPred2A ^65^, using the “long” search mode with default parameters. We took residues with a value > 0.4 to be disordered. We used Anchor from the same IUPred2A package, to find regions within disordered regions that likely harbor linear motifs, using the default Anchor parameters and taking residues with a score > 0.4 to be part of motif-containing regions. For each of these regions (TM, non-TM, ordered, disordered, and motif-embedding disordered regions), we have computed the fraction of non-synonymous mutations that belongs to each mutation category (beneficial, neutral, deleterious and lethal). We then compared these to the respective fractions of the four categories in non-synonymous mutations across the entire polyprotein. We used a one-sided Fisher exact test to test for enrichment (or depletion) in each of the biophysically-defined regions, in comparison with the entire polyprotein, and adjusted the p-values using the Benjamini-Hochberg ^66^ correction. Fig 6 shows the significance of these tests. We plot the relative enrichment for different categories of mutations across different biophysical regions. Relative enrichment is computed as the difference between the fraction of occurrence in the tested region and the fraction of occurrence in the entire polyprotein, divided by the occurrence in the entire polyprotein. For example, relative enrichment of lethal mutations in TM region is calculated as: 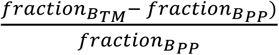

### Cross viral strain and species analysis

We have aligned and compared the conservation of each residue in the polyprotein of the dengue strain we used (strain 2) with DENV1, 3, and 4 serotypes (REF) using CLUSTAL W ^67^. We extracted from the multi-sequence alignment the residues that are conserved across the four species (identical), residues that are substituted by a similar residue, and residues that have dissimilar substitutions or gaps. We then compared the distribution of mutations from the four categories, based on our experimental data analysis (beneficial, neutral, deleterious, and lethal mutations) with their distribution across the entire polyprotein. This was carried out as described in “**Biophysical analysis**.”

## Legends to Extended Data Figures

**Extended Data Figure 1**: Phenotypic characterization of passaged viral populations. (a) Quantification of virus production, quantified by focus forming assay for both replicates. (b) Intracellular RNA content determined by qRT PCR. (c) Efficiency of Plating (EOP) data represented in the embedding in Fig. 1g. EOP was determined based on comparison to the adapted cell line.

**Extended Data Figure 2:** Genotypic characterization of passaged DENV2 populations. (a) PCA loadings of genetic diversity in sequenced populations. The first and second components are host specific, while the third and fourth capture replicate specific differences between the populations. (b) PC scores of individual allele variants. Each score represents the contribution of the allele to the specific pattern of variance captured in the component. Major alleles are highlighted.

**Extended Data Figure 3**: (a) Box plots of the nine individual mutation rate estimates obtained for each passaged population, indicating transitions (“Ts”) and Transversions (“Tv”). (b) Distribution of mutational fitness effects for all possible alleles in the population. The bars in the histogram are shaded to show the proportion of alleles called as Beneficial, Neutral, Deleterious or Lethal, according to their 95% CI. (c) Filled Histogram showing the sequencing depth required to observe alleles of a given fitness class for all populations. (d) UpSet Plots ^68^ comparing shared alleles of individual fitness classes between experimental sets. These comparisons reveal the stochastic nature of beneficial mutations, which are largely unique to the individual populations. Deleterious and lethal mutations act more deterministically, and have more universal effects on fitness across the different host environments.

**Extended Data Figure 4:** (a) *Fitness Wave* representations of the allele fitness dynamics of all of the experimental populations. (b) Line plot showing the change in expected fitness of a randomly drawn allele in the population over time. (c) Line plots showing the mean number of mutations per genome of each fitness class in the sampled genomes used to estimate population fitness.

**Extended Data Movie 1:** Animation of the allele frequencies in the adapting populations over nine passages. Colors: Orange, non-synonymous mutations; Green, synonymous mutation; and Grey, mutations in the UTR.

**Extended Data**: All data has been deposited and is available at the persistent URL: https://purl.stanford.edu/gv159td5450

**Extended Data Table 1:** Table of fitness estimates, mutation classifications, and computational statistics for all data presented here.

**Extended Data Table 2:** Fitness and disorder prediction, and TM prediction data used as input for analysis in Fig 6.

**Extended Data Table 3:** Statistical analyses for enrichment data presented in Fig. 6.

**Extended Data Table 4:** Class-specific mutation rate and error estimates.

## ACKNOWLEDGEMENTS

Research reported in this publication was supported by National Institutes of Health grants AI127447 (JF), AI36178, AI40085, AI091575 (RA), F32GM113483 (PTD), a DARPA Prophecy Award and fellowships from the Naito Foundation (ST) and Uehara Memorial Foundation (ST). We thank the Frydman and Andino labs for discussions and Prof. Marc Feldman and Dmitri Petrov and their labs for constructive comments on the work.

