## Supplementary figures and images for "Principles of dengue virus evolvability derived from genotype-fitness maps in human and mosquito cells"

### Extended Data 1

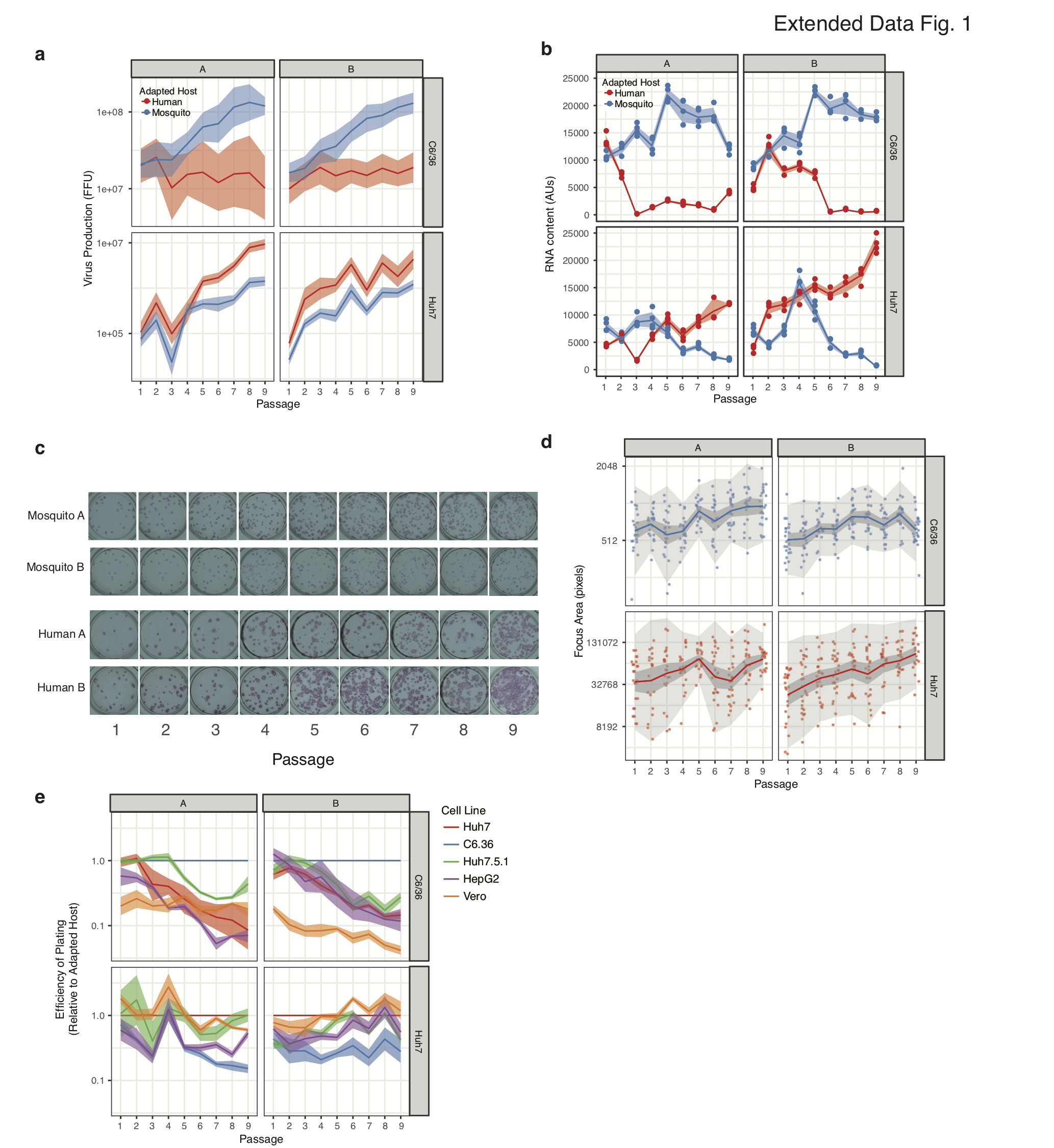

### Extended Data 2

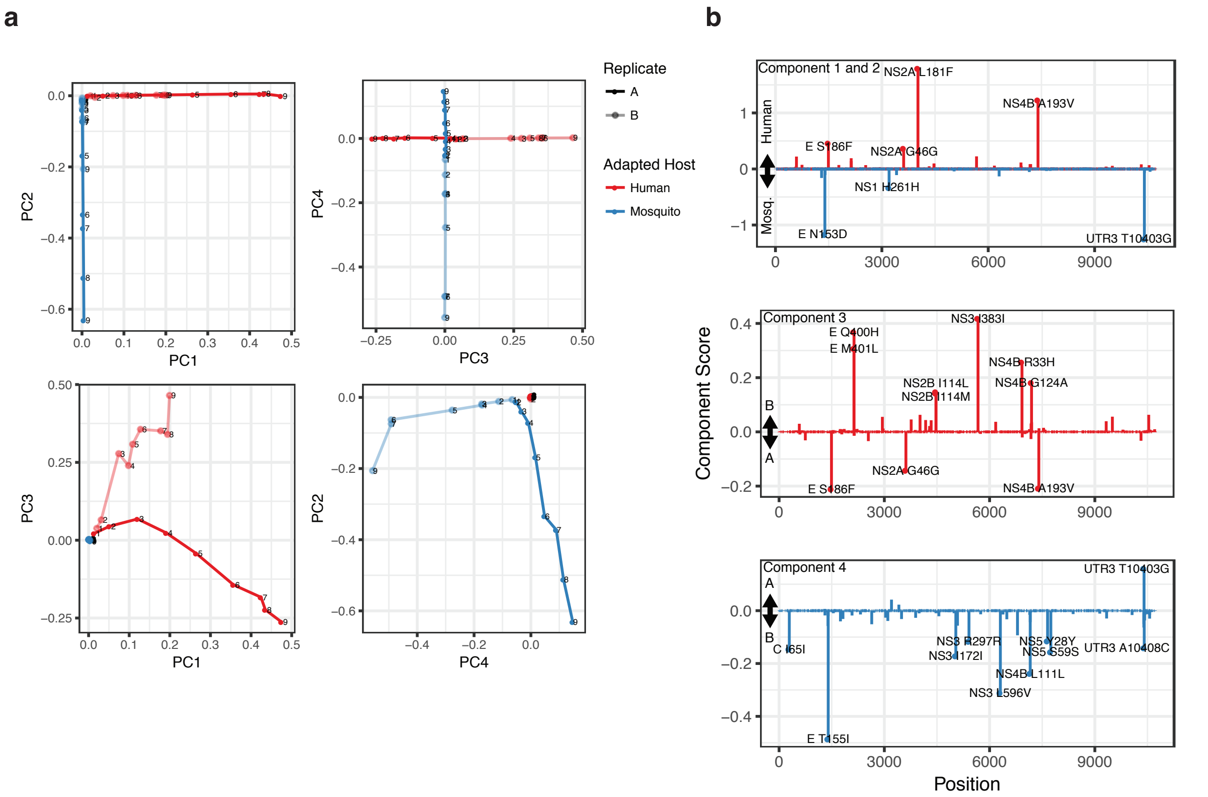

### Extended Data 3

Extended Data Fig. 3

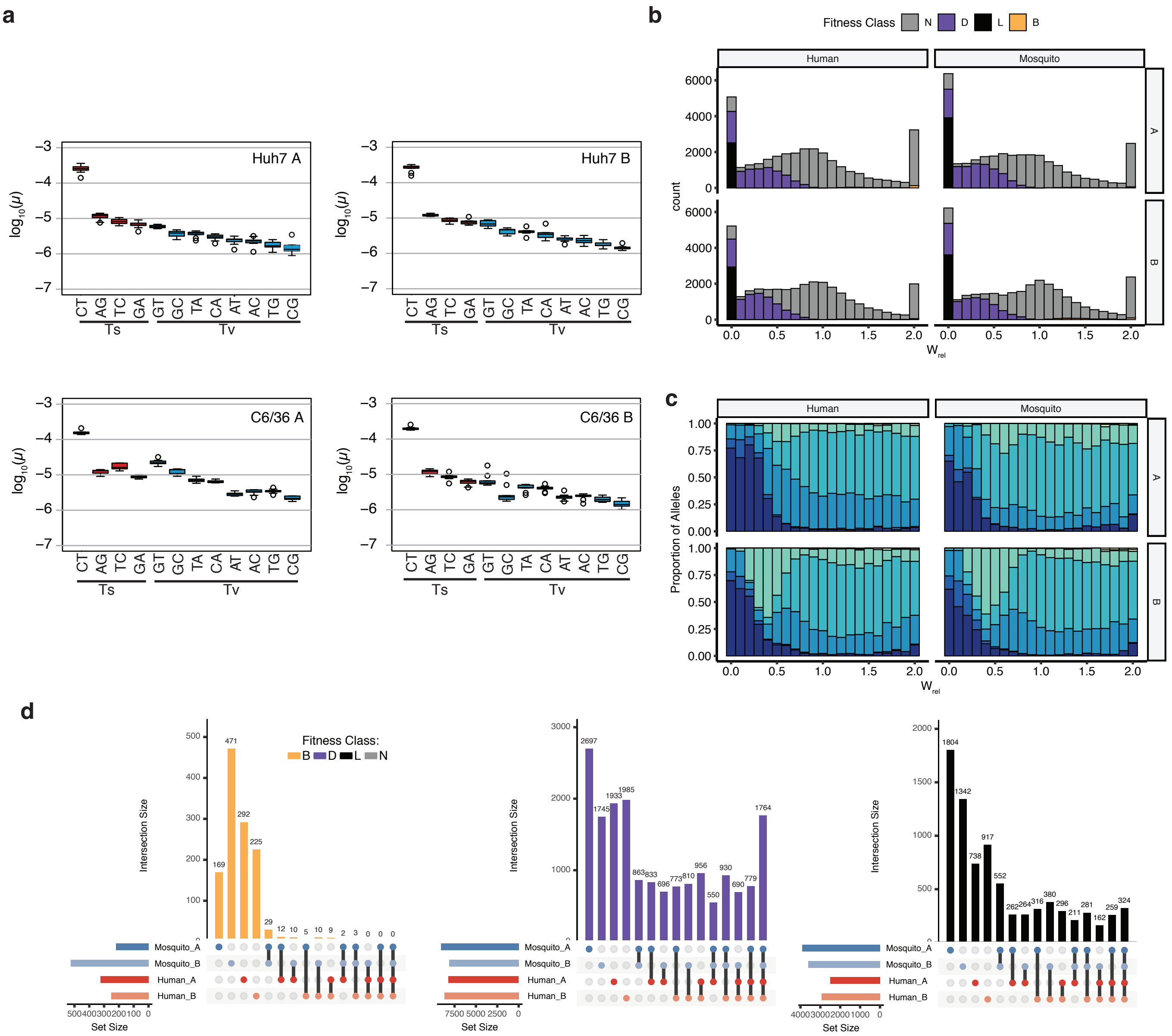

### Extended Data 4

Extended Data Fig. 4

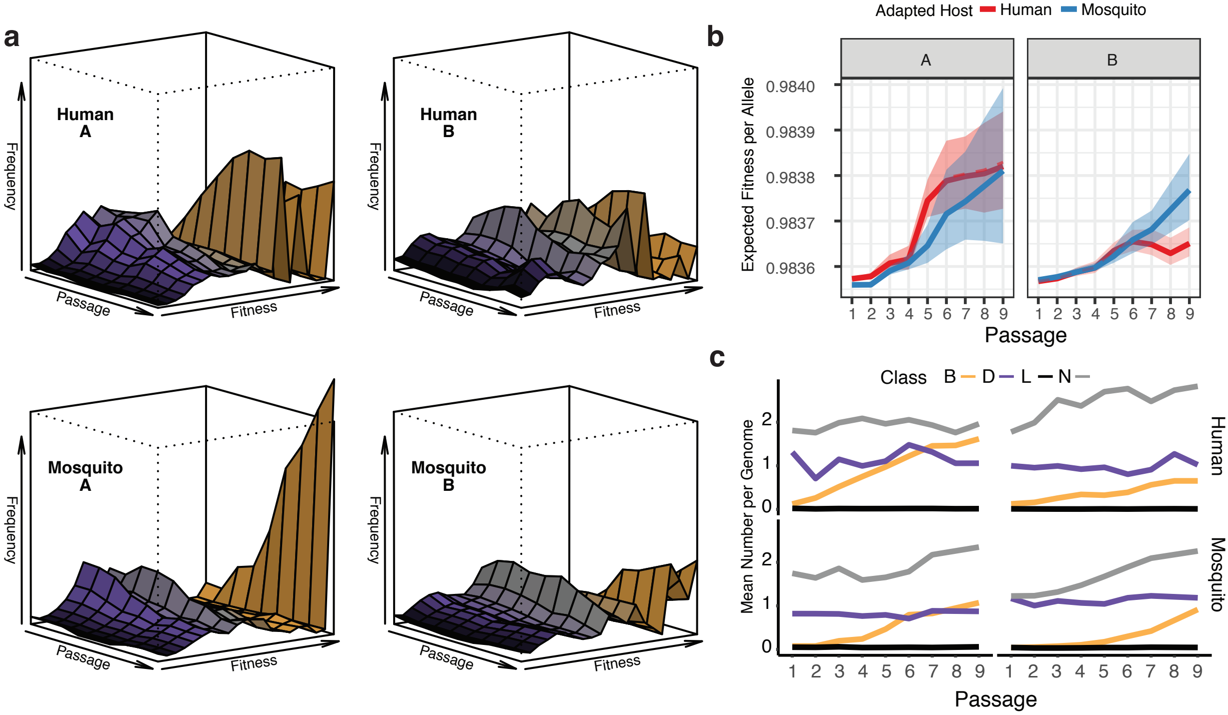
